# DNA double strand break position leads to distinct gene expression changes and regulates VSG switching pathway choice

**DOI:** 10.1101/2021.07.15.452463

**Authors:** Alix Thivole, Ann-Kathrin Mehnert, Eliane Tihon, Emilia McLaughlin, Annick Dujeancourt-Henry, Lucy Glover

## Abstract

Antigenic variation is an immune evasion strategy used by *Trypanosoma brucei* that results in the periodic exchange of the surface protein coat. Underlying this process is the movement of variant surface glycoprotein genes in or out of a specialized locus known as bloodstream form expression site by homologous recombination, facilitated by blocks of repetitive sequence known as the 70-bp repeats, that provide homology for gene conversion events. DNA double strand breaks are potent drivers of antigenic variation, however where these breaks must fall to elicit a switch is not well understood. To understand how the position of a break influences antigenic variation we established a series of cell lines to study the effect of an I-SceI meganuclease break in the active expression site. We found that a DNA break within repetitive regions is not productive for VSG switching, and show that the break position leads to a distinct gene expression profile and DNA repair response which dictates how antigenic variation proceeds in African trypanosomes.

## INTRODUCTION

*Trypanosoma brucei* exists as an extracellular parasite in the mammalian host, colonizing the blood, adipose tissue and skin^1,2^. Survival in the bloodstream is dependent on the parasite’s ability to periodically exchange the expressed surface antigen, the variant surface glycoprotein (VSG), for one that is immunologically distinct^3^. A large proportion of the trypanosome subtelomeric nuclear genome encodes for *VSG* genes and their associated sequences, providing a vast repertoire of potential surface antigens for immune evasion^4^. The *VSG* is expressed from a subtelomeric locus called a bloodstream form expression site (BES) by RNA Polymerase I at an extra nucleolar transcription factory termed the expression site body (ESB)^5,6^ which also recruits the VSG exclusion (VEX) complex^7–9^. BESs are polycistronic transcription units which contain several protein coding genes termed expression site associated genes (*ESAGs*) along with the *VSG*. Although there are approximately fifteen BESs in the nuclear genome^10^, monoallelic expression of a single BES ensures that only one VSG is exposed to the immune system at a time.

The BESs are fragile, and DNA double strand breaks (DSB) in the active BES trigger VSG switching by gene conversion, using the subtelomeric *VSG* archive for repair^11–13^. *VSG* genes are flanked by stretches of 70-bp repeats upstream which serve as recombination substrates for repair^14^, and promote access to the archival *VSGs*^12^ allowing for a greater *VSG* diversity to be expressed during the course of an infection. Antigenic variation appears to follow a semi-predictable order in the selection and expression of *VSG* donors for recombination during the course of an infection^15–18^. The predominant mechanism for switching is via recombination-based *VSG* gene conversion (GC) events, where the expressed *VSG* is replaced with a silent *VSG* gene. Less frequently switching occurs via *in situ* switching which involves the concomitant silencing of the active ES and activation of a silent ES, without any DNA rearrangements^19^. Homologous recombination (HR) is crucial for successful antigenic variation and several factors have been implicated in its success. Factors including BRCA2^20^, RAD51, the RAD51-paralogs^21,22^ and the MRN (MRE11-RAD50-NBS1) complex^23^ have been shown to be required for both efficient HR and antigenic variation. Depletion of the RNases H1 and H2, RTR complex (RECQ2-TOPO3a-RMI1) and the RECQ2 helicase leads to increased *VSG* gene switching by recombination^24,25^. Conversely, ATR kinase and a histone acetyltransferase HAT3, suppress recombination driven VSG switching^26,27^, reinforcing the intimate link between HR and antigenic variation. Disruption of BES chromatin integrity additionally leads to increased DNA accessibility across the BES and switching of the active BES by HR^28–34^. However, the molecular machinery that facilitates VSG switching remains elusive.

VSG switch events are triggered by DSBs in the BES^11,13^ however, little is known about how the position of the DSB influences antigenic variation or if breaks within the 70-bp repeats trigger VSG switching. Although the 70-bp repeats are dispensable^35^, they have been shown to direct DNA pairing and provide homology during switching^13,36^. We sought to further understand how the position of the DSB in the active BES determined the VSG switching outcome. DSBs in the active BES, either adjacent to the telomere, immediately upstream of the active *VSG* or adjacent to the *VSG* promoter have been shown to be highly toxic, but VSG switching is only triggered when it falls immediately upstream of the active *VSG*^13,37^. Equally, naturally occurring DSBs have been detected in both the active and silent BESs, including the 70-bp repeats of the active BES^13,37^. This led to the intriguing hypothesis that the nature of the 70-bp repeats rendered them fragile and prone to DSBs, therefore triggering antigenic variation. But this has not been experimentally tested. Here, we have assessed the relationship between the position of a DSB in the active BES, changes in genes expression immediately following a DSB and the repair outcome. We have found that when a DSB in the active BES leads to a significant loss of VSG transcript, VSG switching is triggered by homologous recombination.

## MATERIALS AND METHODS

### *Trypanosoma brucei* growth and manipulation

Lister 427, MITat1.2 (clone 221a), bloodstream stage cells were cultured in HMI-11 medium^38^ at 37.4 °C with 5 % CO_2_. Cell density was determined using a haemocytometer. For transformation, 2.5 x 10^7^ cells were spun for 10 minutes at 1000g at room temperature and the supernatant discarded. The cell pellet was resuspended in prewarmed cytomix solution^39^ with 10 μg linearised DNA and placed in a 0.2 cm gap cuvette, and nucleofected (Lonza) using the X-001 program. The transfected cells were placed into one 25 cm^2^ culture flask per transfection with 36 ml warmed HMI-11 medium only and place in an incubator to allow the cells to recover for approximately 6 hours. After 6 hours, the media was distributed into 48-well plates with the appropriate drug selection. Strains expressing TetR and Sce ORF and with an I-SceI recognition-sites upstream of the active *VSG*-ESs (VSG^up^)^11^ have been described previously. G418, and blasticidin were selected at 2 μg.ml^-1^ and 10 μg.ml^-1^ respectively. Puromycin, phleomycin, G418, hygromycin, blasticidin and tetracycline were maintained at 1 μg.ml^-1^. Clonogenic assays were plated out with 32 cells per plate under non-inducting conditions for all cell lines and either 32 cells per plate or 480 cells per plate under inducing conditions. Plates were counted 5-6 days later and subclones selected for further analysis. Puromycin sensitivity was assessed using 2 μg.ml^-1^.

### Cell line set up

To construct p70int^sce^, we synthesised pMA-RQ-70int (Invitrogen) which contains a 315 bp block of 70-bp repeats with a XcmI site embedded within the repeats, 150 bp from either end. pMA-RQ-70int was linearized with Xcm1 (New England Biolabs) and dephosphorylated with Calf Intestinal Phosphotase (MBI Fermantas) to introduce the *RFP^sce^PAC*^40^. pARDR^s^P was digested with Xcm1 digestion (NEB) to release the *RFP^sce^PAC* and ligated into the linearized pMA-RQ-70int. Before transfection p70int^sce^ was linearized with Not1 (NEB).

To construct pESAG1^sce^, pMA-T-ESAG1 was synthesised (Invitrogen) to contain 300 bp of sequence homologous to the region downstream of *ESAG1* with and XcmI site 150 bp from either end. The *RFP^sce^PAC* was integrated as with p70int^sce^ and linearised by NotI. To generate the Pseudo^sce^ cell line we used a long primer approach. Using primers PseudoRFPF ( CAACAAAATTATAGCAGAATGCAACGTCGACAAAAGGCTCAAGAAATTAACGGCCTACAC GCGGGTCCCATTGTTTGCCTCT) and PseudoPACR ( GTTTTGGCGCGTTGTTCCGTATCTGCTGAGCAAACCTTTTGCGCCGGCTGCTGCGGCGGATAACTATTTTCTTTGATGAAAG) we amplified the *RFP^sce^PAC* cassette using Phusion Polymerase (ThermoFisher). 5 μg of PCR product was used for transfection using standard protocols. Plasmids used to generate a double knock-out of the *RAD51* gene and *MRE11* are described in^23,37^.

### Immunofluorescence microscopy

Immunofluorescence analysis was carried out using standard protocols as described previously^37^. Rabbit anti-VSG2 was used at 1:20 000 and rabbit anti-γH2A^41^ was used at 1:250. Fluorescein-conjugated goat α-rabbit and goat anti-mouse secondary antibodies (Pierce) were used at 1:2000.

Samples were mounted in Vectashield (Vector Laboratories) containing 4, 6-diamidino-2-phenylindole (DAPI). In *T. brucei*, DAPI-stained nuclear and mitochondrial DNA can be used as cytological markers for cell cycle stage^42^; one nucleus and one kinetoplast (1N:1K) indicate G_1_, one nucleus and an elongated kinetoplast (1N:eK) indicate S phase, one nucleus and two kinetoplasts (1N:2K) indicate G2/M and two nuclei and two kinetoplasts (2N:2K) indicate post-mitosis. Images were captured using a ZEISS Imager 72 epifluorescence microscope with an Axiocam 506 mono camera and images were processed in ImageJ.

### RNA Analysis

RNA samples were taken at 0 and 6 hours post I-SceI induction. RNA was extracted from 50 ml of culture at 1 x 10^6^ cells / ml. RNA-seq was carried out on a BGISeq platform at The Beijing Genome Institute (BGI). Reads were mapped to a hybrid genome assembly consisting of the T. brucei 427 reference genome plus the bloodstream VSG-ESs^10,33,43^. Bowtie 2-mapping was used with the parameters --very-sensitive --no-discordant --phred33. Alignment files were manipulated with SAMtools^44^. Per-gene read counts were derived using the Artemis genome browser^45^; MapQ, 0. Read counts were normalised using edgeR and differential expression was determined with classic edgeR. RPKM values were derived from normalised read counts in edgeR^46^.

### DNA analysis

Genomic DNA was extracted from 50 ml of a >1 x 10^6^ cells-ml culture, spun at 1000 g for 10 minutes and the pellet was resuspended in 200 μl of PBS. Genomic DNA was prepared using the DNeasy Tissue Kit (Qiagen). For Southern Blot analysis, purified DNA was digested with I-SceI (NEB) and AvrII (NEB) and run on a 0.75 % agarose gel at 75V until there was adequate separation. The gels were washed in 0.25M HCl for 15 minutes followed by two 10 minutes washes in dH_2_O. The DNA was denatured for 45 minutes (1.5 M NaCl (Sigma), 0.5 M NaOH.), followed by two washes in neutralization solution (1.5 M NaCl (Sigma), 0.5 M Tris base (Sigma), p.H. to 7.0 with HCl) for 45 minutes and a final wash in in 2x SSC (3 M NaCl (Sigma), 0.3 M sodium citrate (Sigma), p.H. to 7.0 with HCl). The DNA was transferred on a Zeta-probe nylon membrane (Bio-Rad) by capillary action (according to the standard protocol) overnight. The DNA was cross-linked to the membrane by UV irradiation on a Stratalinker (Stratagene). The membrane was washed in a pre-hybridization solution (6 x SSC, 1 x Denhardt’s, 100 μg/ml denatured Herring Sperm DNA (Sigma) and 0.5 % SDS) at 65°C for 2 hours, and incubated overnight in hybridization containing the ^32^P probes at 65°C. Before to exposing the membrane to a phosphor screen the membrane was washed twice in a washing solution (0.2 x SSC and 0.2 % SDS). The phosphor screen was exposed for at least 48 hours before visualization on a Phosphor-imager (Amersham). The *PAC* probe was a 618 bp EcoRI fragment of the pPac plasmid. The VSG221 probe was a PCR product using primers VSG221F and VSG221 R (details below).

For PCR assays, PCR fragments with lengths is above 500 bp to 2 kb were carried out with GoTaq enzyme (Promega) according to standard protocols. Analysis of subclones was previously described^11,26,47^ and used the following primers VSG221F (CTTCCAATCAGGAGGC), VSG221R (CGGCGACAACTGCAG), RFP (ATGGTGCGCTCCTCCAAGAAC), PAC (TCAGGCACCGGGCTTGC), ESAG1F (AATGGAAGAGCAAACTGATAGGTTGG), ESAG1R (GGCGGCCACTCCATTGTCTG), Pseudo F (GTACGCGGCCGCTGCCTCTAGCAGTTGCGCCG) and Pseudo R (GTACCATATGTTAATTAATGCATCCATCTTTGTATTCC). To validate correct integration of the RFP:PAC cassette the following primers were used: PseudoFval (CGCTATTCGGAACAGGAAAG) PseudoRval (CACCCCAGGCTGTTGTAAGT) for Pseudo^sce^ and ESAG1F and PseudoR for ESAG1^sce^. PCR across the 70-bp repeats were carried out with LongAmp enzyme (NEB) according to standard protocols with primers PseudoF and PACR or RFPF and VSG221R.

### qPCR and RT-qPCR analysis

To determine the dynamics of cleavage we followed a published pPCR protocol^25^. Briefly, 1 x 10^6^ cells were collected at 3 hours intervals between 0-24 hours post I-SceI induction. Genomic DNA was extracted using the Qiagen Blood and Tissue kit and quantified using a Nanodrop. The DNA was diluted to 0.2 ng.μl^-1^ and 1 ng was analysed by qPCR using Luna Universal qPCR MasterMix (NEB) with 500nM of primers. For each pair of primers (below), triplicates of each sample were run per plate (Hard-shell PCR Plates 96 well, thin wall; Bio-Rad), which were sealed with Microseal ‘B’ Seals (BioRad). All experiments were run on a CFX96 Touch Real-time Detection system with a C1000 Touch Thermal cycler (Bio-Rad), using the following PCR cycling conditions: 95°C for 3 min, followed by 40 cycles of 95°C for 15 sec and 60°C for 1 min (fluorescence intensity data collected at the end of the last step). Data was then analysed by relative quantification using the ΔΔCt method (CFX Maestro software - BioRad). Primer pairs CACAACGAGGACTACACCATC and CGGCCTATTACCCTGTTATCC (ESAG1^sce^, Pseudo^sce^, 70int^sce^), or GTTGTGAGTGTGTGCTTACC and ATCTAGAGGATCTGGGACCC (VSG^up^)^25^. The expression levels of ESAG6/7 and VSG2 following induction of a break were determined using the following protocol. RNA was extracted using a Qiagen RNeasy Kit and the samples were treated with DNase 1 for 1 hour according to manufactures instructions and eluted in 30 μl of RNase free water. The samples were quantified using a Nanodrop (ThermoFisher). cDNA was prepared using SuperScript IV (ThermoFisher) following the supplier instructions from 1-2 μg RNA with a polyT primer. For each pair of primers (used at 500nM), triplicates of each sample were run per plate (Hard-shell PCR Plates 96 well, thin wall; Bio-Rad), which were sealed with Microseal ‘B’ Seals (BioRad). All experiments were run on a CFX96 Touch Real-time Detection system with a C1000 Touch Thermal cycler (Bio-Rad), using the following PCR cycling conditions: 95°C for 3 min, followed by 40 cycles of 95°C for 15 sec and 60°C for 1 min (fluorescence intensity data collected at the end of the last step). Data was then analysed by relative quantification using the ΔΔCt method (CFX Maestro software - Bio-Rad) and Cq determination regression was used. Primer pairs VSG2F (AGCTAGACGACCAACCGAAGG) and VSG2R (CGCTGGTGCCGCTCTCCTTTG)^48^ or ESAG6/7 F (ACTGTGGATGAATTGGCGAA) ESAG6/7 R (ACTGCTACTGTGTTGGACCC). In all cases, product abundance was determined relative to an actin control locus, which was amplified with primer pairs actin F (GTACCACTGGCATTGTTCTCG) and actin R (CTTCATGAGATATTCCGTCAGGTC).

## RESULTS

### A DSB in the 70-bp repeats of the active BES does not lead to VSG switching

Natural DSBs have been detected in both silent and active BESs and within the 70-bp repeats^13,37^, but whether these breaks lead to a productive VSG switching has not been directly tested. To do this we used the established tetracycline-inducible I-SceI meganuclease system^37,49,50^, and integrated an I-SceI recognition site within the major block of 70-bp repeats in BES1, the active BES (Fig 1A; 70int^sce^) and in parallel, selected two additional positions to test in BES1 where the break is in close proximity to 70-bp repeats but distal to the active *VSG*. In addition to the 70int^sce^ cell line we integrated an SceR between *ESAG1* and the smaller block of 70-bp repeats (Fig 1A; ESAG1^sce^), and between two blocks of 70-bp repeats in the *Pseudo* gene (Fig 1A; Pseudo^sce^), as has been previously described^13^. To validate correct integration in the 70int^sce^ cell line, we performed southern blotting and PCR assays (Supplementary Fig 1). Following digestion with I-SceI and AvrII, the parental cell line gives a 5.8 kb fragment, while the 70int^sce^ will give a fragment of 3.9 kb (Supplementary Fig 1A). Additional validation using a PCR assay, confirmed the integration of the *RFP:PAC* cassette in the 70-bp repeats (Supplementary Fig 1A). To validate correct integration in the Pseudo^sce^ and ESAG1^sce^ cell lines we used specific PCR assays (Supplementary Fig 1 B and C). Following validation, we set up survival assays to assess the effect of a DSB in the 70int^sce^, Pseudo^sce^ and ESAG1^sce^ cell lines. As a control for VSG switching, we used a previously published cell line, where the I-SceI recognition site is immediately upstream of the active *VSG* and adjacent to the large block of 70-bp repeats (Fig 1A; VSG^up^), and which shows high rates of VSG switching following induction of a DSB^13,37^. In the VSG^up^ cell line, only 5 % of the cells survive a DNA break, which agrees with previously published data^23,25,37^ (Fig 1B). To our surprise, 50 % of the cells in the ESAG1^sce^ cell line, 75 % in Pseudo^sce^ and 95 % in 70int^sce^ survived the DSB (Fig 1B). This was confirmed when looking at the effect of an I-SceI break only (Fig 1C), here the proportion of survivors was calculated by dividing the number of induced survivors by the number of uninduced survivors. This is in stark contrast to other positions tested in the active BES, where in addition to the VSG^up^ cell line, a DNA break adjacent to the active VSG promoter or at the telomere / chromosome junction resulted in between 20 - 5 % survival respectively^37^. It was only when an SceR site was inserted into a non-transcribed silent BES that more than 80 % cell survival was recorded^37^. A DSB in the *Pseudo* gene of the active BES has been reported before^13^, and in our study served as an additional control, however due to differences in the efficiency of cutting it is not possible to directly compare survival. From our data we conclude that an I-SceI induced DSB that is flanked by 70-bp repeats and distal to the *VSG* gene is not toxic to bloodstream form trypanosomes.

**Figure 1:**
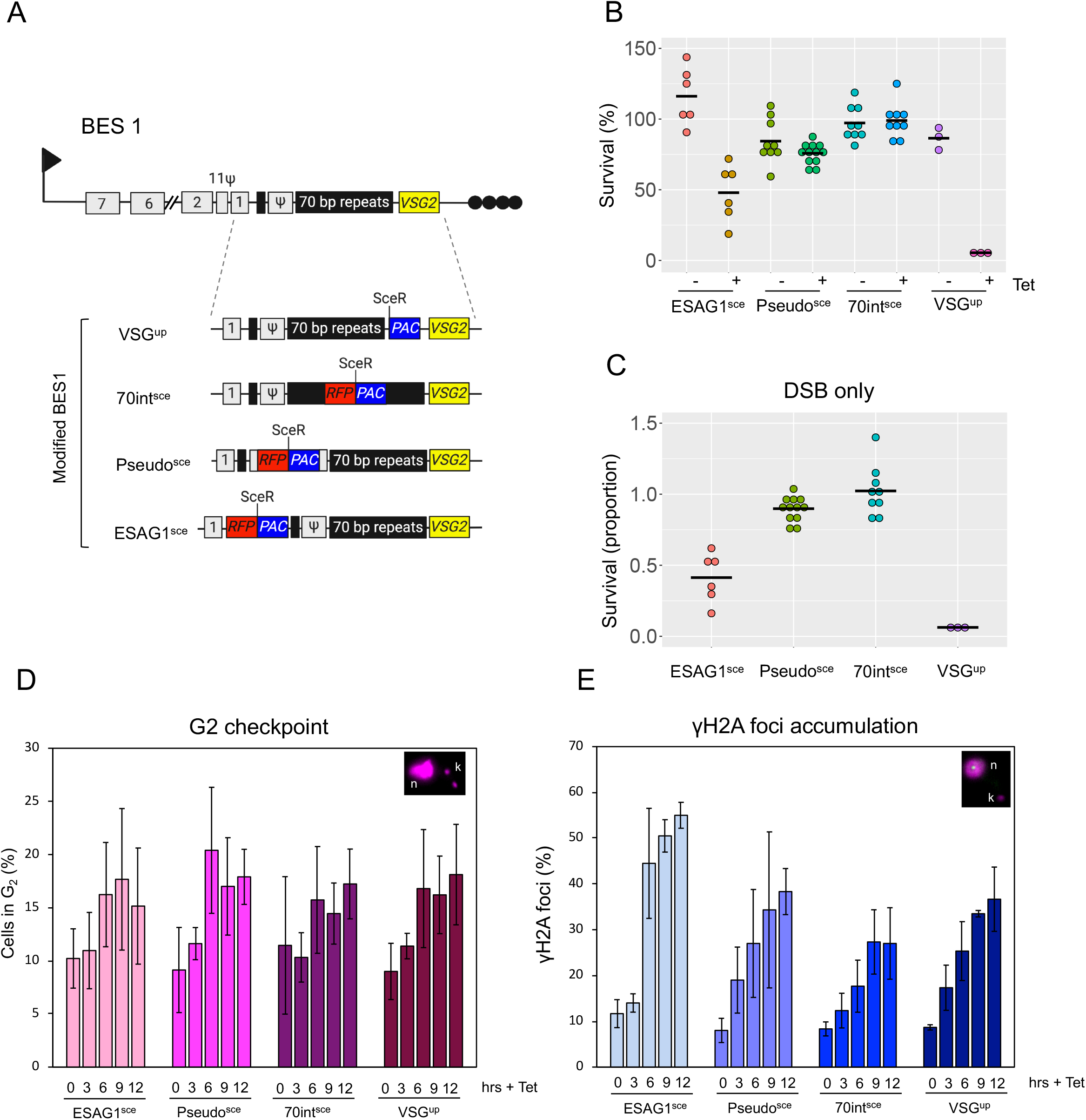
Double strand breaks upstream of the active VSG are not typically lethal. (A) Schematic showing the *VSG2* BES on chromosome 6a with the individual modifications. I-SceI recognition site (SceR) and reporter genes incorporated to give VSG^up^^37^; 70int^sce^; Pseudo^sce^ and ESAG1^sce^. Arrow, BES *RNA Pol1* promoter; grey boxes, ESAGS; black boxes, 70 bp repeats; yellow box, VSG; black circles, telomere. *RFP:PAC, Red Fluorescent Protein and Puromycin N - Actyltransferase and PAC;* black circles, telomere. (B) Clonogenic survival assay, cells were plated out into 96 - well plates in either non or I-SceI inducing conditions. Clones were counted after 7 days. Black bar represents the mean. (C) Proportion of survivors following a I-SceI break. Number of plates are as follows: VSG^up^ – 3 per condition; 70int^sce^ – 9 per condition; Pseudo^sce^ – 9 for the – Tet and 12 for the + Tet and ESAG1^sce^ – 6 per condition. (D) Cell in G2 were determined by DAPI staining and scored according to the position of the nucleus (n) and kinetoplast (k) following an induction of an I-SceI break. n=2 (independent inductions,) 200 cell counted for each time point. (E) Focal accumulation of γH2A as assessed by immunofluorescence assay following an induction of an I-SceI break. n=2 (independent inductions,) 200 cell counted for each time point.

Using the G2 cell cycle checkpoint and γH2A focal accumulation^11,41^ to assess the DNA damage response following a DSB in the BES, we found that the number of cells in G2 and the number of nuclei with γH2A foci increased in all four cell lines between 0 - 12 hours (Fig 1D and E). In the ESAG1^sce^, Pseudo^sce^, 70int^sce^ and VSG^up^ cells lines we saw a 1.7 (9 hours), 2.2 (6 hours), 1.3 (12 hours) and 1.8 (12 hours) fold increase in cells in G2 respectively (Fig 1D). For the γH2A focal accumulation, we saw a 4.7, 4.7, 3.2 and 4.2-fold increase in foci at 12 hours post DSB in the ESAG1^sce^, Pseudo^sce^, 70int^sce^ and VSG^up^ respectively (Fig 1E). We noted that for both cells in G2 and γH2A focal accumulation, the weakest response was seen in the 70int^sce^ cell line.

### Repair of DSBs within or upstream of the 70-bp repeats does not result in BES sequence loss

The high rate of survival following a break in the 70int^sce^, Pseudo^sce^ and ESAG1^sce^ cell lines suggests a mechanism of repair that occurs either in the absence of any significant deficit to the BES, or a reduction in the efficiency of I-SceI cleavage within sequence we are targeting. The presence of both native and exogenous DNA within the BES provides a number of markers that can be used to assess both cleavage and BES reorganization following the DNA break - repair cycle. We used individual subclones (ESAG1^sce^ n=24; Pseudo^sce^ n=25; 70int^sce^ n=25; VSG^up^ n=20) to establish how the survivors repaired the DNA break. To determine the efficiency of cleavage, all clones were tested for puromycin sensitivity. Following a DSB, resection results in loss of *RFP:PAC* ORF. Sensitivity to puromycin is therefore used as an indicator for cutting by I-SceI while puromycin resistance suggests no cutting (Fig 1A). Of the clones tested, one ESAG1^sce^ clone and one 70int^sce^ clone were found to be puromycin resistant (Fig 2A). All the other clones were puromycin sensitive. This suggests that cleavage by I-SceI is equally efficient across all sites tested. In the ESAG1^sce^, Pseudo^sce^, 70int^sce^ cell lines, all the clonal survivors were positive for VSG2 by immunofluorescence assay (Fig 2B) indicating none had undergone VSG switching. The results from the Pseudo^sce^ clones broadly agrees with previously published data^13^, where a DSB in this position has been shown to exhibit a low switching frequency^12^. The slight discrepancy between observed VSG switching may be due to technical variations between the two studies. Clonal survivors from the VSG^up^ cell line were assessed in parallel as a control (n=20) and were all VSG2 negative as previously published (Fig 2B)^13,37^. These results suggest that a DSB in or upstream of the 70-bp repeats are poor catalysts for VSG switching.

**Figure 2:**
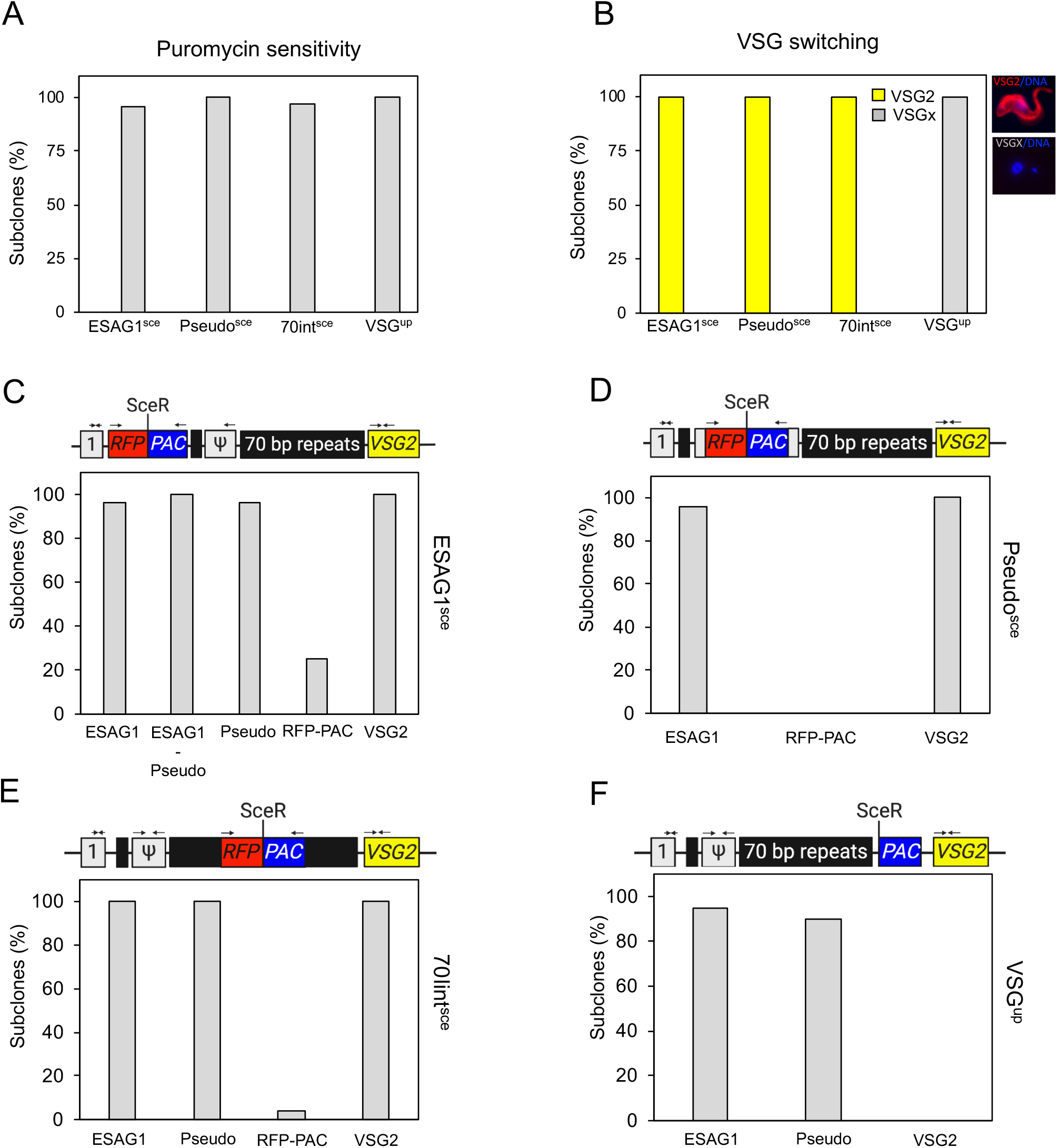
A DSB in or upstream of the 70-bp repeats does not lead to VSG switching. (A) Induced clonal survivors were assessed for sensitivity to puromycin. (B) Induced clonal survivors were assessed and scored for VSG2 expression by immunofluorescence assay. (C) ESAG1^sce^ clonal survivors were assessed by PCR assay to demonstrate the presence or absence of *ESAG1, Pseudo, RFP:PAC* and *VSG2* within the modified BES. n= 25 (D) Pseudo^sce^ clonal survivors were assessed by PCR assay to demonstrate the presence or absence of *ESAG1, RFP:PAC* and *VSG2* within the modified BES. n= 25 (E) 70int^sce^ clonal survivors were assessed by PCR assay to demonstrate the presence or absence of *ESAG1, Pseudo, RFP:PAC* and *VSG2* within the modified BES. n= 25 (F) VSG^up^ clonal survivors were assessed by PCR assay to demonstrate the presence of absence or *ESAG1, Pseudo* and *VSG2* within the modified BES. n= 20. See the schematic maps in Supplementary Figure 2–5 for details of the primer sites. grey box ‘1’, ESAG1; grey box Ψ, pseudo gene; black boxes, 70 bp repeats; yellow box, VSG2; *RFP:PAC, Red Fluorescent Protein and Puromycin N – Actyltransferase*.

We next assessed the DNA rearrangements following repair in the cloned survivors. Using primers specific to genes in BES1 or the *RFP:PAC* cassette we used their presence or absence to infer the mechanisms of repair. In the ESAG1^sce^ cell line, all clones, except clone 2, tested had retained *ESAG1, pseudo* and *VSG2* (Fig 2C and Supplementary Fig 2). The presence of the *VSG2* gene was expected as the cells were positive for VSG2 by immunofluorescence (Fig 2B). Given that 23 out of the 24 clones were puromycin sensitive we expected to see loss of the *RFP:PAC* cassette, however, 3 clones (clones 5, 7, 14) amplified a product that was reduced in size (Supplementary Fig 2A). The full-length *RFP:PAC* cassette was amplified in the single puromycin resistant clone - clone 17, as expected given it is uncut (Supplementary Fig 2). To determine the extent of sequence loss in the remaining 20 clones we used primers to amplify a product from *ESAG1* - *Pseudogene* (Supplementary Fig 3B). The puromycin resistant clones (clone 23; Supplementary Fig 2A and 2B) and the parental cell line gave PCR fragments approximately 6 kb - the size expected if the *RFP:PAC* cassette is intact. The remaining *RFP:PAC* negative clones gave PCR fragments of approximately 5 kb, equivalent to that of wild-type cells suggesting these clones had repaired back to the wild-type state (Supplementary Fig 2B). The PCR products were then sequenced and in 13 clones we could detect microhomology mediated end joining (MMEJ) scars, where short sequences of homology are used to align and join the ends. In 5 we could not detect any MMEJ scar, suggesting repair by HR.

We then assessed the Pseudo^sce^ and 70int^sce^ clones. In both cell lines, all the clones had retained *ESAG1* and *VSG2* but lost the *RFP:PAC* (Fig 2D and E and Supplementary Fig 3 and 4), apart from the single 70int^sce^ puromycin resistant clone (clone 10) that had retained the *RFP:PAC* cassette as expected (Fig 2E and Supplementary Fig 4). These results agree with the immunofluorescence showing VSG2 on the surface of all the survivors (Fig 2B). In VSG^up^ cell line, all the clones had lost *VSG2*, 18 had retained the *Pseudo* gene and 19 had retained *ESAG1* (Fig 2F and Supplementary Fig 5) which agrees with previously published data^23,37^.

### DSB repair in the 70-bp repeats is independent of RAD51 and MRE11

In trypanosomes, RAD51 is required for HR and plays a crucial role in antigenic variation^11,22,51,52^ as does MRE11 which is also required for processing of single strand (ss) DNA at a BES^23,53,54^. We generated RAD51 null mutants in the ESAG1^sce^, Pseudo^sce^, 70int^sce^ cell lines (Supplementary Fig 6) and MRE11 null mutants in the 70int^sce^ cell line (Supplementary Fig 7A). In the *rad51* nulls the survival was reduced by approximately 80 % in the ESAG1^sce^, 60 % in the Pseudo^sce^ and 80 % in the 70int^sce^ cell lines (Fig 3A). The reduction in survival is in line with what has been reported previously for *rad51* nulls^11^. We next looked at the cloning efficiency following induction of an I-SceI DSB. By comparing the cloning efficiency, we see that the majority of repair in the 70int^sce^*rad51* cell line is RAD51-independent (proportion of survival 1 vs 0.8 Fig 1C and Fig 3B). In both the ESAG1^sce^*rad51* and Pseudo^sce^*rad51* null cell lines, survival was reduced compared to the 70int^sce^*rad51* but there was no significant difference compared to the respective parental cell lines (ESAG1^sce^ vs ESAG1^sce^*rad51* - 0.4 vs 0.3 (Fig 1C and Fig 3B); Pseudo^sce^ vs Pseudo^sce^*rad51* - 0.9 vs 0.5 (Fig 1C and Fig 3B)). Additionally, we studied the impact of MRE11 in the 70int^sce^ cell line. We generated MRE11 nulls in this cell line to determine if repair by MRE11 dependent MMEJ dominated at this position. MRE11 has been implicated in repair by MMEJ in eukaryotes^55^ but not in Leishmania^56^. In the 70int^sce^*mre11* null cells only 40 % of the uninduced population was able to grow (Fig 3A). Following an I-SceI break, approximately 35 % of the cells were able to survive a DSB (survival proportion of 0.9; Fig 3A and B) and PCR assays revealed that the cells had lost the *RFP:PAC* cassette but retained *VSG2* and *ESAG1*, as has the parental 70int^sce^ cell line, this suggests that repair at this position is MRE11-independent (Supplementary Fig 7B).

**Figure 3:**
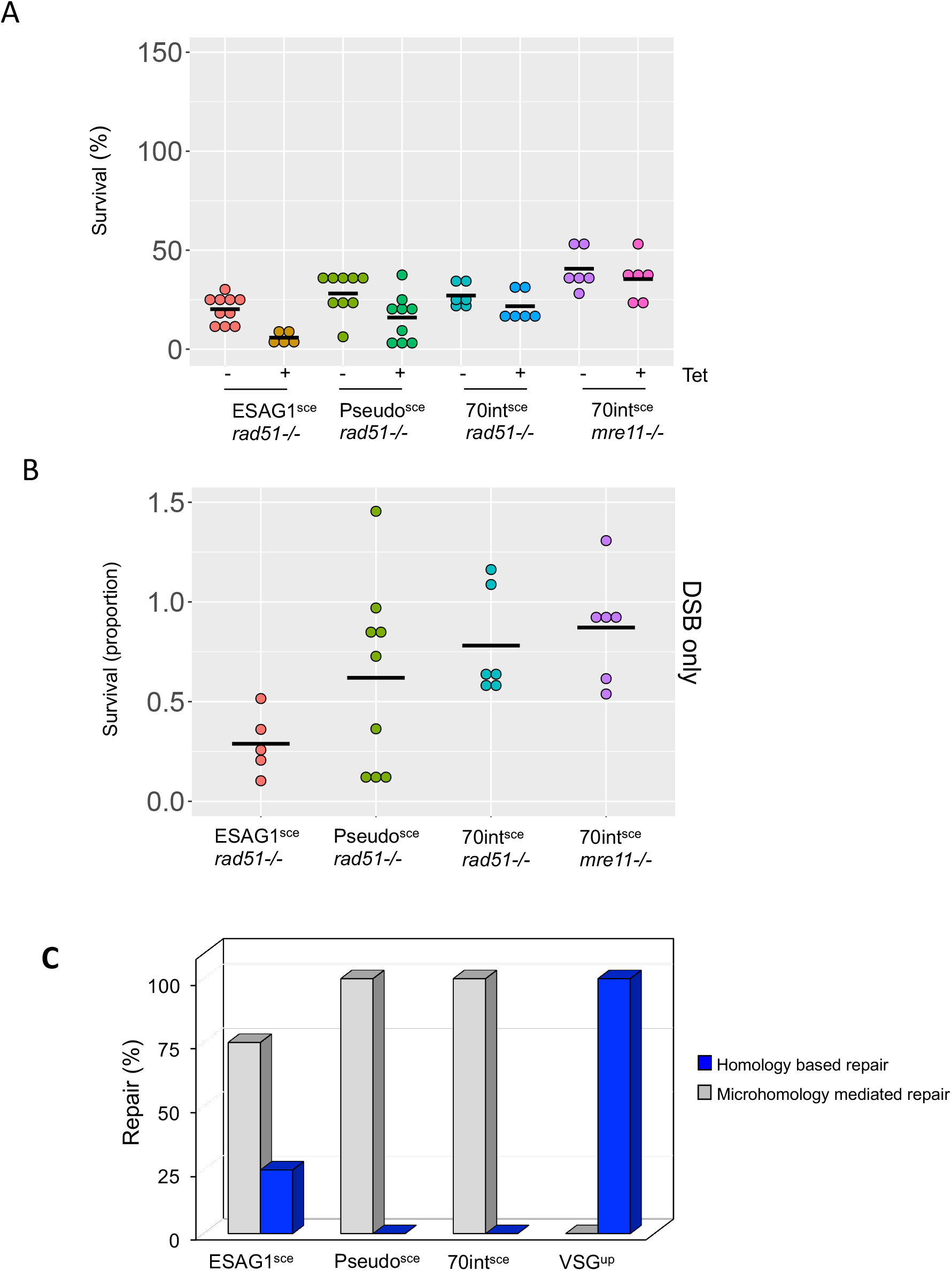
Repair within the 70-bp repeats is independent of RAD51 and MRE11. (A) Clonogenic survival assay, cells were plated out into 96 – well plates in either non or I-SceI inducing conditions. Clones were counted after 7 days. Black bar represents the mean. Number of plates for each cell line are as follows: 70int^sce^*rad51^-/-^* and 70int^sce^*mre11^-/-^* 6 plates for each condition; Pseudo^sce^*rad51^-/-^* 9 plates for each condition and ESAG1^sce^ *rad51^-/-^* 9 plates for each condition.(B) Proportion of survivors following a I-SceI break. (C) Percentage of repair pathway choice in clonal survivors.

### The cleavage-repair cycle and not the timing of the DSB determines the VSG switching outcome

The striking difference between DSB repair mechanism and the VSG switching outcome at different positions led us to ask whether these differences could have arisen due to variability in the timing of I-SceI cleavage. We used a qPCR assay previously described^25^ to assess cleavage in all four cell lines. Genomic DNA was prepared between 0 – 24 hours post induction, taking samples every 3 hours (Fig 4A) and a qPCR assay was run using primers that spanned the I-SceI recognition site. The amount of product was reduced in all cell lines as early as 3 hours following a DSB, consistent with previous reports, but most dramatically in the ESAG1^sce^ cell line, where cutting appears almost complete by three hours. In both the ESAG1^sce^ and Pseudo^sce^ cell line, no product was detected at 24 hours suggesting I-SceI cleavage was complete (Fig 4A). In the VSG^up^ and 70int^sce^ cell lines, the SceR site was cleaved in more than 60 % of the cells after 24 hours. Although we did not detect an increase in the amount of PCR product in the VSG^up^ cell line as seen in Devlin, *et al*^25^, we do detect a slight increase between 9 - 12 hours (Fig 4A, black line), the data are broadly consistent between the two studies. We next looked at *VSG2* expression by RT-qPCR^48^. RNA was extracted at 6-hour intervals following I-SceI induction, and as anticipated, VSG2 expression decreased over the time course in the VSG^up^ cell line, closely matching that of I-SceI cleavage, with a 50 % reduction in *VSG2* expression at 18 hours (Fig 4A and B). In the ESAG1^sce^ cell line rapid cutting resulted in an initial drop in *VSG2* expression, which returned close to baseline level at 18 hours. A similar pattern was observed in the 70int^sce^ cell line where although more than 50 % of the cells had undergone I-SceI cleavage the reduction in level of VSG2 expression returned over 80 % at 18 hours (Fig 4A and B). The *VSG2* RT-qPCR revealed an increase in *VSG2* transcript in the Pseudo^sce^ cell line, peaking at 6 hours post DSB, which again returned wild-type levels at 18 hours (Fig 4B). These results suggest that the activated repair mechanism and subsequent VSG switching outcome were not influenced by the timing of I-SceI cleavage, but rather whether or not *VSG* transcription was inhibited.

**Figure 4:**
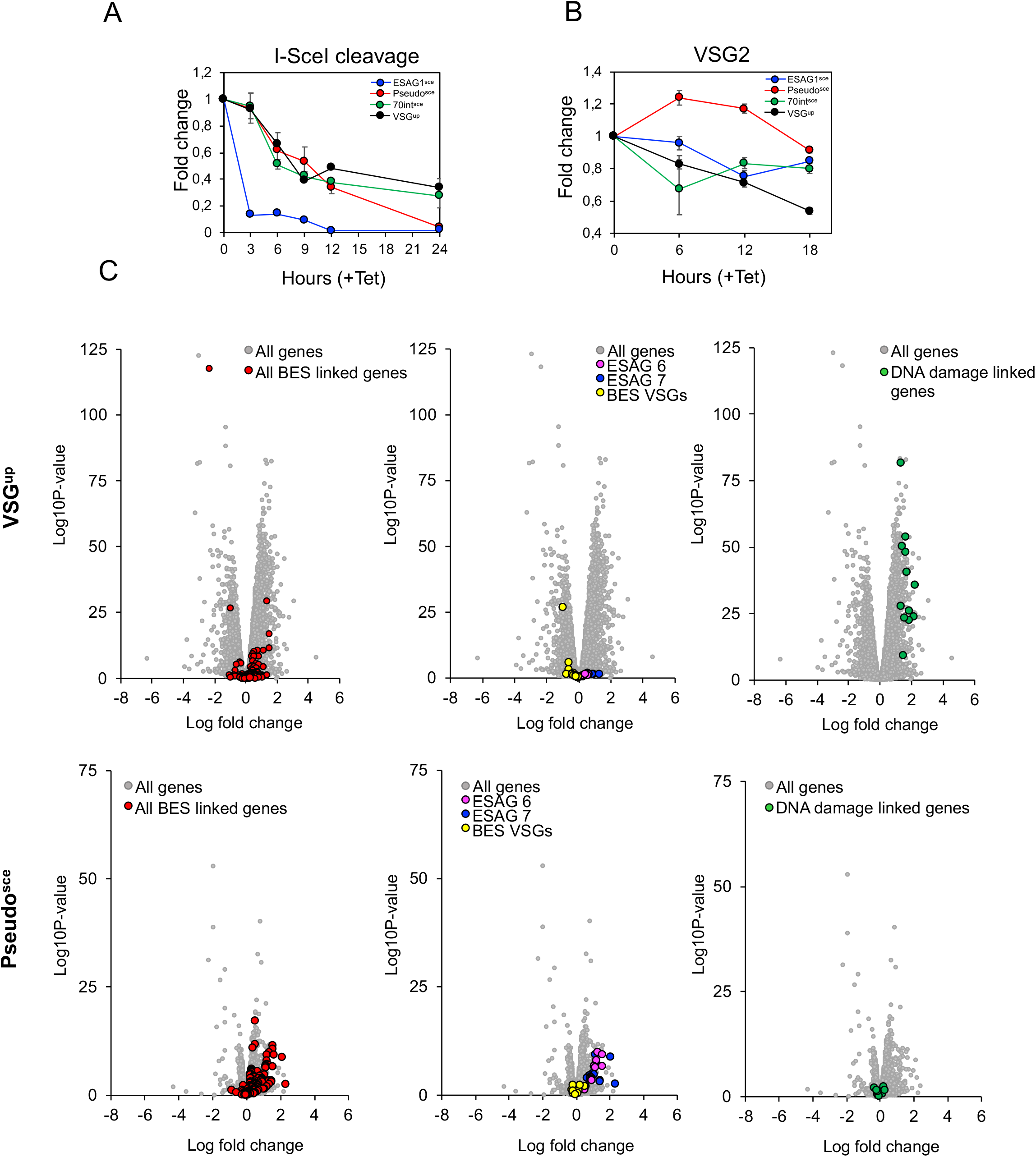
A DNA break at the active BES elicits a specific change in gene expression. (A) qPCR assay to assess I-SceI cleavage following induction. (B) RT-qPCR analysis of the expression levels of *VSG2* at 6-hour intervals following an I-SceI DNA break. (C) RNA-seq analysis at 6 hours following a DSB in the VSG^up^ cell line and the Pseudo^sce^ cell line. Values are averages of three independent replicates relative to wild-type controls. Red circles BES linked genes; yellow circles BES VSGs; pink circles, BES *ESAG6*; blue circles, BES *ESAG7*; green circles, DNA damage linked genes (All genes are included in Table 1).

### The position of a DSB in the active BES leads to distinct gene expression changes

To explore gene expression changes following a DSB we analysed the transcriptome in the Pseudo^sce^ and the VSG^up^ cell line – comparing a cell line that exhibits 100 % VSG switching versus one where no VSG switching was detected following a DSB (Fig 4C and Supplementary Fig 8; Table 1). The increase in *VSG2* expression in the Pseudo^sce^ cell line led us to look first at BES linked genes. In the VSG^up^ cell line, we noted that there were very few changes in the BES linked genes following a break (Fig 4C upper panel: VSG^up^ and Fig 5). In comparison, although the Pseudo^sce^ cell line showed a more constrained change in the gene expression profile following a break than the VSG^up^ cell line, a specific cohort of BES linked genes was upregulated in response to a break at 6 hours (Fig 4C lower panel: Pseudo^sce^). Closer analysis of this cohort of genes revealed that BES linked ESAG7 and ESAG6 are specifically upregulated in the Pseudo^sce^ cell line (16 x *ESAG7* genes FC 0.48 to 2.3 and Log_10_ 1.64 to 10.64; 14 x *ESAG6* FC 1.62 to 0.46 and Log_10_ 0.95 to 11,32) (Fig 4C). Assessment of all BES-linked genes (i.e., from all the silent BESs and the active BES) using a generic BES - which follows the overall conservation in gene order from BES promoter to VSG^57^, to compare gene expression changes revealed that *ESAG7* (FC of 0.345 vs 1.224; *P value = <0.0001*) and *ESAG6* (FC of 0.339 vs 1.100; *P value = <0.0001*) were significantly upregulated 6 hours following a DSB in the Pseudo^sce^ cell line as well as *ESAG 3Ψ* (FC of 0.211 vs 0.767; *P value = 0.0471*) and *ESAG 8* (FC of 0.066 vs 0.536; *P value = <0.0001*) (Fig 5A). The VSG^up^ cell line showed a significant increase in expression in *ESAG 4* (FC of 0.678 vs 0.219; *P value = 0.03*) only. We then assessed whether there were any changes in the expression of the BES linked *VSG* genes (14 in total for both cell lines) and observed a small change in the Pseudo^sce^ cell line (Fig 5A) only. Looking specifically at changes in gene expression across the active BES, a similar pattern was seen with the Pseudo^sce^ cell line, *ESAG7* and *ESAG6* were upregulated (FC 0.58 vs 1.56 and 0.4 5 vs 1.6) (Fig 5B) and in the VSG^up^ cell line, *ESAG 5Ψ* and ESAG 4 were upregulated (FC 0.29 vs 1.39 and 0.39 vs 0.15, respectively) (Fig 5B). Surprisingly, the biggest change seen in the active BES, was a 2 - fold reduction in expression of the *Pseudo* gene in the VSG^up^ cell line (Fig 5B).

**Figure 5:**
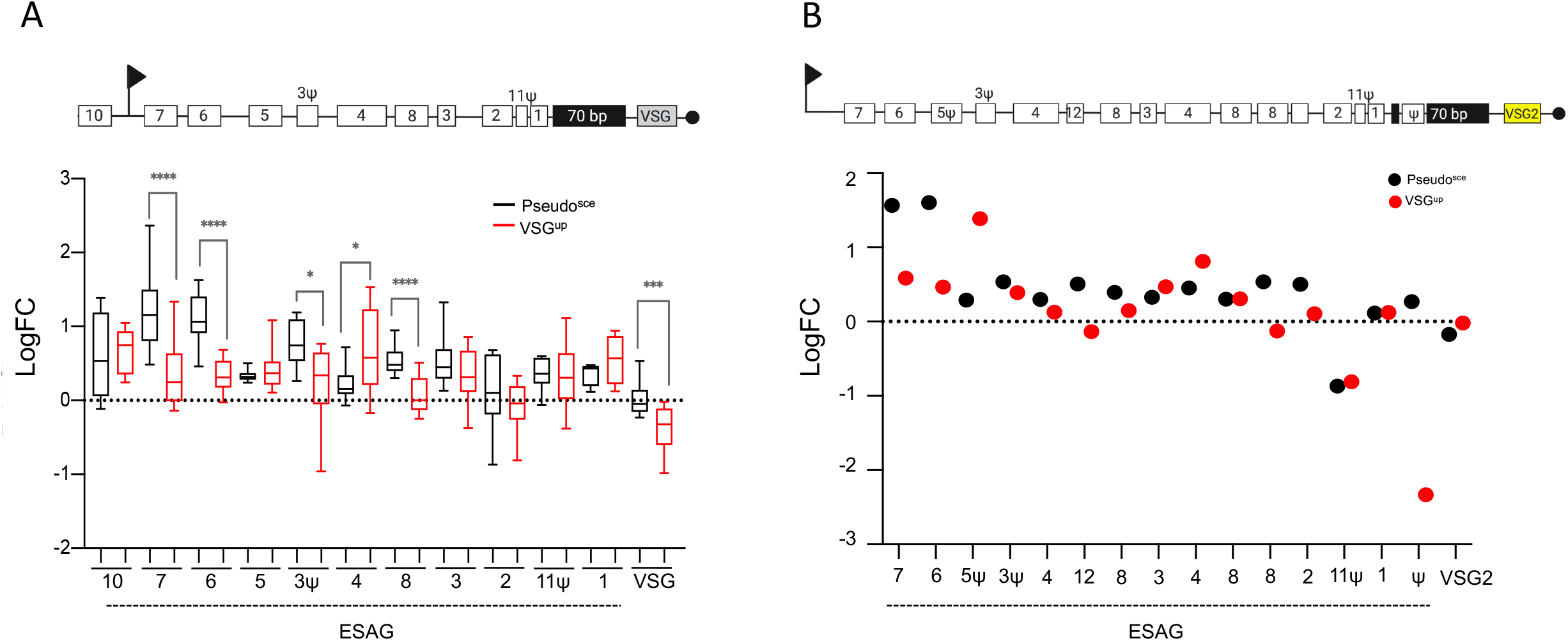
A DSB in the active *pseudo* gene leads to an increase in expression in *ESAG6* and *7*. (A) Upper panel: Schematic depicting a generic BES. Lower panel: The box-plot depicts the sum of BES linked ESAG and VSG genes expression; whiskers extend from the minimal to the maximal value. Horizontal line indicates the mean. (B) Upper panel: Schematic depicting the actively expressed BES1 with *VSG2* at the telomeric end. Lower panel: The plot depicts the sum of BES1 linked ESAG genes and VSG2 expression. (All genes are included in Table 1).

To determine, which, if any, genes were upregulated following a DNA break, we took an arbitrary cutoff of the top 100 genes with the greatest fold changes in both the Pseudo^sce^ and the VSG^up^ cells lines. As already reported here, in the Pseudo^sce^ cell line this list was dominated by *ESAG6* and *7* (Table 2 - transferrin receptor). We observed that for the VSG^up^ cells line, 22 genes were ‘hypothetical conserved’, which is not surprising given that over 60 % of the trypanosome genome is unannotated (Table 2) and of the remaining genes several of those that were annotated could be associated with a role in DNA damage and repair (Fig 4C). These include NBS1 which forms part of the MRN complex which has been shown to promote repair by HR in trypanosomes^23^. Here we see an upregulation in the expression of NBS1 (Tb927.8.5710; FC 1.67 and Log_10_ 53,56). RAD5 is required for post replication repair^58^ and here we see an upregulation in RAD5 expression (Tb927.1.1090; 1.86 FC and Log_10_ 22.19). Two putative helicase genes were upregulated Tb927.6.920; FC 1.86 and Log_10_ 25.88 and Tb927.9.13610; FC 1.71 and Log_10_ 40.15), and the role of DNA helicases in DNA repair are well described^59^. Protein kinases play a key role in the DNA damage response, coordinating the cellular response that preserves genome integrity^60^. Four kinase genes were upregulated, a MEKK-related kinase 1 (Tb927.10.14300; FC 1.33 and Log_10_ 27.52), a CMGC/SRPK protein kinase (Tb927.7.960; FC 1.63 and Log_10_ 47.94) - which was also identified in a genome-wide screen for DNA damage in trypanosomes^61^, a serine/threonine-protein kinase (Tb927.10.1910; FC 1.40 and Log_10_ 49.9) and a inositol hexakisphosphate kinases (Tb927.8.3410; FC 2.28 and Log_10_ 35.5), which have been shown to regulate DNA repair via conserved signalling pathways that sense damage^62^. SUMOylation and ubiquitination are both implicated in DNA repair^63^ and within this cohort of genes, two proteins were annotated as SUMO-interacting motif containing proteins (Tb927.3.1660 and Tb927.8.1290) and one as a ubiquitin-conjugating enzyme (Tb927.3720). SUMOylation is also required for VSG expression in trypanosomes^48^. These results show that as early as 6 hours the position of a DSB can lead to specific gene expression changes that position the cell to either invoke a repair pathway that leads to VSG switching or partially activate silent BES promoters as a backup should repair damage the integrity of the BES.

## Discussion

DSBs in the active expression site, although highly toxic, are a potent driver of antigenic variation^13,37^. What leads to the formation of these DSBs is unknown, but it is unlikely to be a process directed by an endonuclease, and rather a function of the location of the BES in the subtelomeres or a characteristic of the BES itself^64^. In this study we show that a DSB in repetitive regions in the BES are poor triggers for antigenic variation, and that where MMEJ dominates as the major form of repair, silent BES promoters are partially activated as a means to facilitate rapid *in situ* switching should repair be deleterious to the cell. We also show that the position of the break leads to distinct gene expression changes and regulates repair pathway choice and the VSG switching outcome.

### DNA breaks in repetitive sequence do not trigger antigenic variation

Found within the BES and associated with silent *VSG* genes in the subtelomeric arrays, the 70-bp repeats have been shown to facilitate HR^12,35^. It has been hypothesized that the nature of the repetitive sequence may lead to break formation thereby acting as the trigger for VSG switching. By establishing a series of cell lines, we were able directly assess the effect of a DSB embedded within or flanked by repetitive sequence, in the active BES. We then assessed the DNA damage response and repair pathway choice in these cell lines and individual clones. Our results indicate that a DSB within the central portion of the 70-bp repeats or upstream - in the *Pseudo* gene or adjacent to *ESAG1*, do not display the same toxicity as elsewhere in the active BES^37^, and are not productive for antigenic variation. In the three positions tested, we do note that all trigger the DNA damage response as indicated by an increase in γH2A foci, and cells in stalled in G2 position in the cell cycle, but the response in dampened in the 70int^sce^ cell line. Our data also suggests that the majority of repair in the 70-bp repeats or upstream happens via RAD51 and MRE11 independent MMEJ. In contrast a DSB proximal to the *VSG* is repaired by homology-based mechanisms as previously shown^37^. In mammalian cells, MRE11 is critical MMEJ initiation^65^, but both in Trypanosomes^23^ and the related kinetoplast parasite *Leishmania*^56^, MMEJ has been shown to be MRE11 independent. We propose that a rapid cleavage - repair cycle facilitated by MMEJ within the 70-bp repeats or in the *Pseudo* gene, where there are an abundance of sequence microhomologies, maintains *VSG* expression, BES integrity and high levels of cell survival. Previous work has shown that a DSB proximal to the active BES promoter^37^ is also repaired by MMEJ without resulting in VSG switching, but with high levels of cell death. We suggest that like in the ESAG1^sce^ cell line, where 50 % of the cells are able to repair and survive, limited microhomologies at these positions result in a lower proportion of survival.

In BES1 – the active BES in our strain, the 70-bp repeats are approximately 5 kb, so it remains possible that breaks at the immediate *VSG* proximal end, but still within the 70-bp repeats, may lead to VSG switching if resection extends into the *VSG* gene or upstream sequence. Upstream of the BES linked *VSG* genes is the co-transposed region (CTR), deletion of the CTR and the 5’ end of *VSG2* leads to rapid switching^66^. We tentatively suggest that for a DNA break to be productive in terms of VSG switching, processing by resection would have to disrupt the CTR or *VSG* expression. Consistently, the only position that shows high frequency VSG switching is following a break adjacent to the 70-bp repeats and up stream of the *VSG*^13,37^.

### DNA break site in the active BES gives rise to distinct gene expression changes that determine VSG switching pathway

Our data, for the first time, shows that there are distinct gene expression changes associated with the position of a DNA break. I-SceI access is broadly equivalent across all sites tested, suggesting the differences in the response to the break are not due to variations in the efficiency of cutting due to the location of the break. Our results indicate that there are distinct gene expression changes as early as 6 hours following a DSB in the active BES and propose that the position of the DSB triggers specific VSG switching pathways which may act to preserve the integrity of the BES. Specifically, we see expression of BES promoter proximal *ESAG 7* and *ESAG 6* in the Pseudo^sce^ cell line, suggesting partial derepression of silent promoters. MMEJ is a mutagenic repair mechanism which results in deletions flanking the site of the break^67^, however it appears to be the favoured form of repair in the active BES (DNA break at the active BES promoter^37^, *ESAG1* gene, *Pseudo* gene and within the 70 bp repeats). Given its mutagenetic potential, in order to prevent loss of *VSG* transcription, or a deletion of a large section of the BES, silent BES promoters are partially activated, pre-adapting the cells to *in situ* switching should they need to in order to survive. In contrast, where a DSB leads to repair by HR and VSG switching, the silent BES promoters are not activated, and we report upregulation of genes required for DNA damage and repair. We note that the FC are small in these analyses, however these samples were taken 6 hours following a DSB. At this early time point only a proportion of the cells had been cut (Fig 4A, 50 % cut in VSG^up^ and 40 % cut in Pseudo^sce^).

It has been previously reported that chemically induced nuclear DNA damage leads to upregulation of silent BES promoter and other RNA polymerase 1 transcribed genes^68^. Our data following a break in the Pseudo^sce^ cell line broadly agrees with this, however we do not see expression of other RNA polymerase 1 genes such as EP1 Procyclin (Tb927.10.10260). This may be due to the specificity of the DSB in this study as compared to broad chemical damage in Sheader et al. Reversible derepression of silent BES promoters is also seen when the active BES promoter transcription is blocked^69^, in a proposed probing of silent BESs before the cells commit to a switch. Our data suggest that this derepression phenotype is rapid, here within 6 hours of a break and specific where a DNA break in the active BES will trigger repair by MMEJ.

### Conclusion

The mechanisms underlying recombination-based VSG switching are not well understood. To our knowledge, this is the first report that demonstrates that the position of a DSB in the active BES can trigger distinct gene expression changes, directing the cell through a specific repair pathway. We report that a DSB in the 70-bp repeats or upstream is rapidly repaired, predominantly by MMEJ, whereas only a DSB within close proximity to the expressed *VSG* leads to recombination-based VSG switching. This data has now expanded our understanding on DSBs as a trigger for VSG switching.

## Data availability

All RNA-seq data has been deposited onto the ENA under the accession number is: PRJEB44245 and unique name: ena-STUDY-INSTITUT PASTEUR-12-04-2021-14:30:14:655-830.

## Acknowledgements

We would like to thank Sebastian Hutchinson for help with the RNA-seq analysis. AKM was supported by the Erasmus + program of the European Union; work in the LG laboratory has received financial support from the Institut Pasteur (G5 Junior group) and the National Research Agency [ANR - VSGREG]. EJM is part of the Pasteur - Paris University (PPU) International PhD Program. This project has received funding from the European Union’s Horizon 2020 research and innovation programme under the Marie Sklodowska-Curie grant agreement No 665807 and from the Foundation Recherche Médicale grant number FDT202012010602. Funding for open access charge: Institut Pasteur G5 funding.

## Author contributions

AT, ADH, AKM, EJM and LG performed the experiments and data analysis. ET performed the RNA-Seq analysis. AT and LG designed the experiments. LG wrote the manuscript and all authors contributed to editing the final draft.

## Conflict of interest

The authors declare that they have no conflict of interest.

---

**Supplementary Figure 1:**
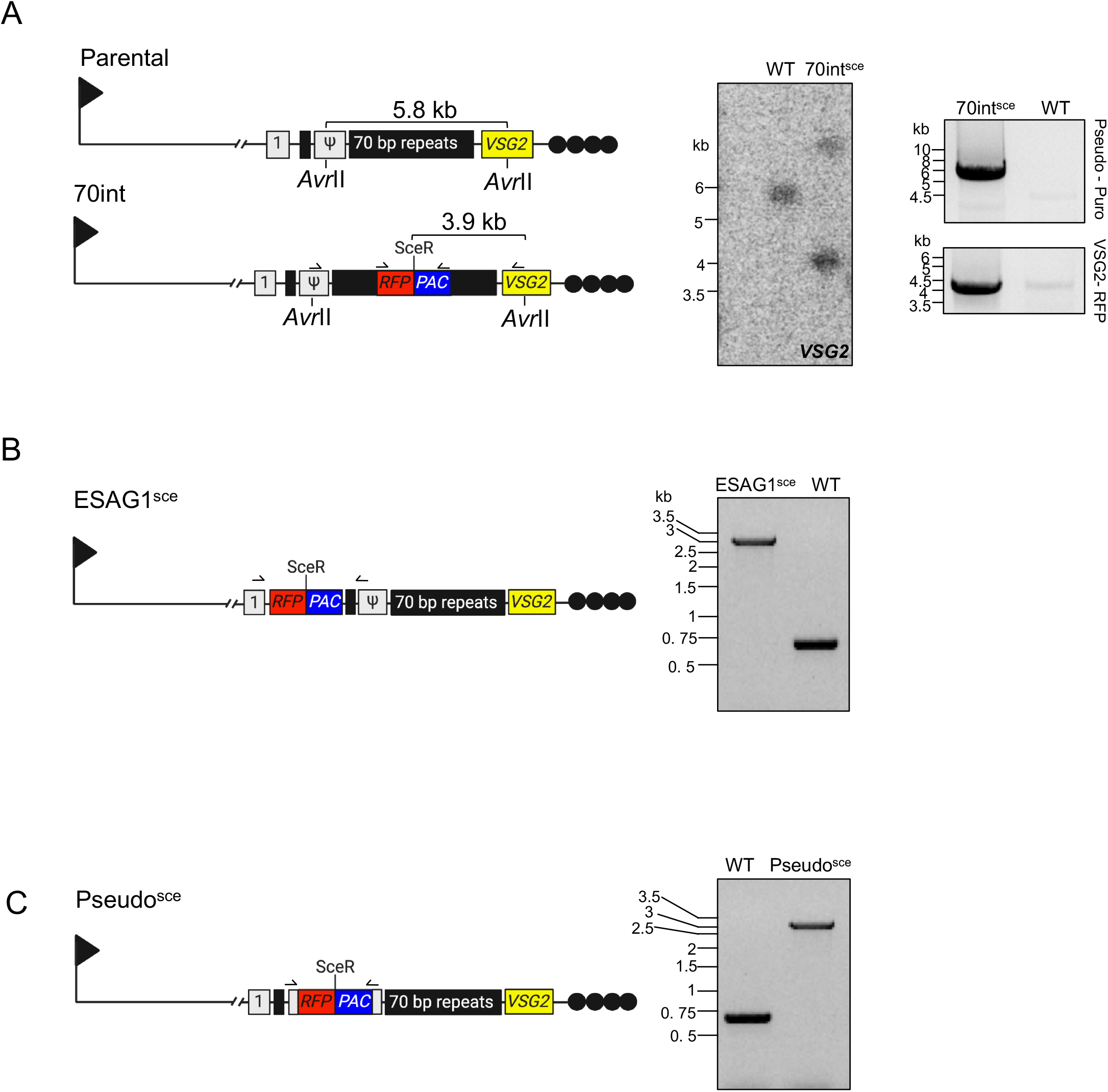
Validation of cell line set up. (A) Left panel: Schematic showing the 70int^sce^ cell line. Middle panel: Southern blot. Right panel: PCR assay, correct integration should give a band of approximately 5000 bp using primers from the pseudo gene to *PAC* and a 4523 bp band from *VSG2* to *RFP*. Relevant restriction sites shown. (B) Left panel: Schematic showing the ESAG1^sce^ cell line. Right panel: PCR assay, correct integration should give a 2334 bp band using primers from *ESAG1* to *PAC*. (C) Left panel: Schematic showing the Pseudo^sce^ cell line. Right panel: PCR assay, wild-type should give a band at 600 bp and correct integration of the *RFP:PAC* cassette should give 2564 bp. Black arrow, BES *RNA Pol1* promoter; grey boxes, ESAGS; black boxes, 70 bp repeats; yellow box, VSG; black circles, telomere. *RFP:PAC, Red Fluorescent Protein and Puromycin N – Actyltransferase and PAC*. Line arrow, primer binding sites.

**Supplementary Figure 2:**
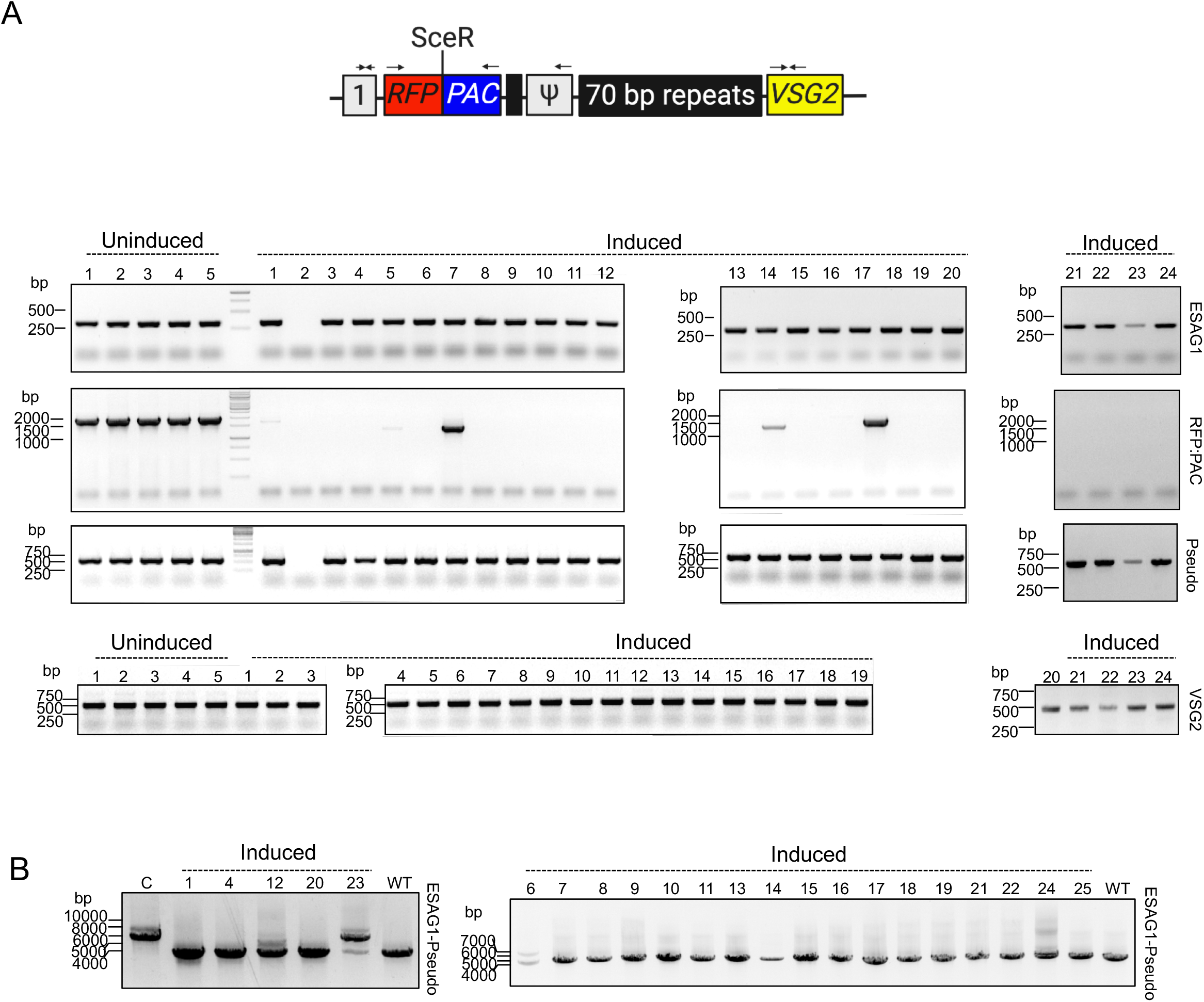
ESAG1^sce^ PCR assays. (A) Schematic indicates the location of the primers used. (B) The PCR assays show the presence or absence of *ESAG1, Pseudo, RFP - PAC* and *VSG2* following an I-SceI induced DSB. (C) PCR assays show the presence or absence of *ESAG1* - *Pseudo* product.

**Supplementary Figure 3:**
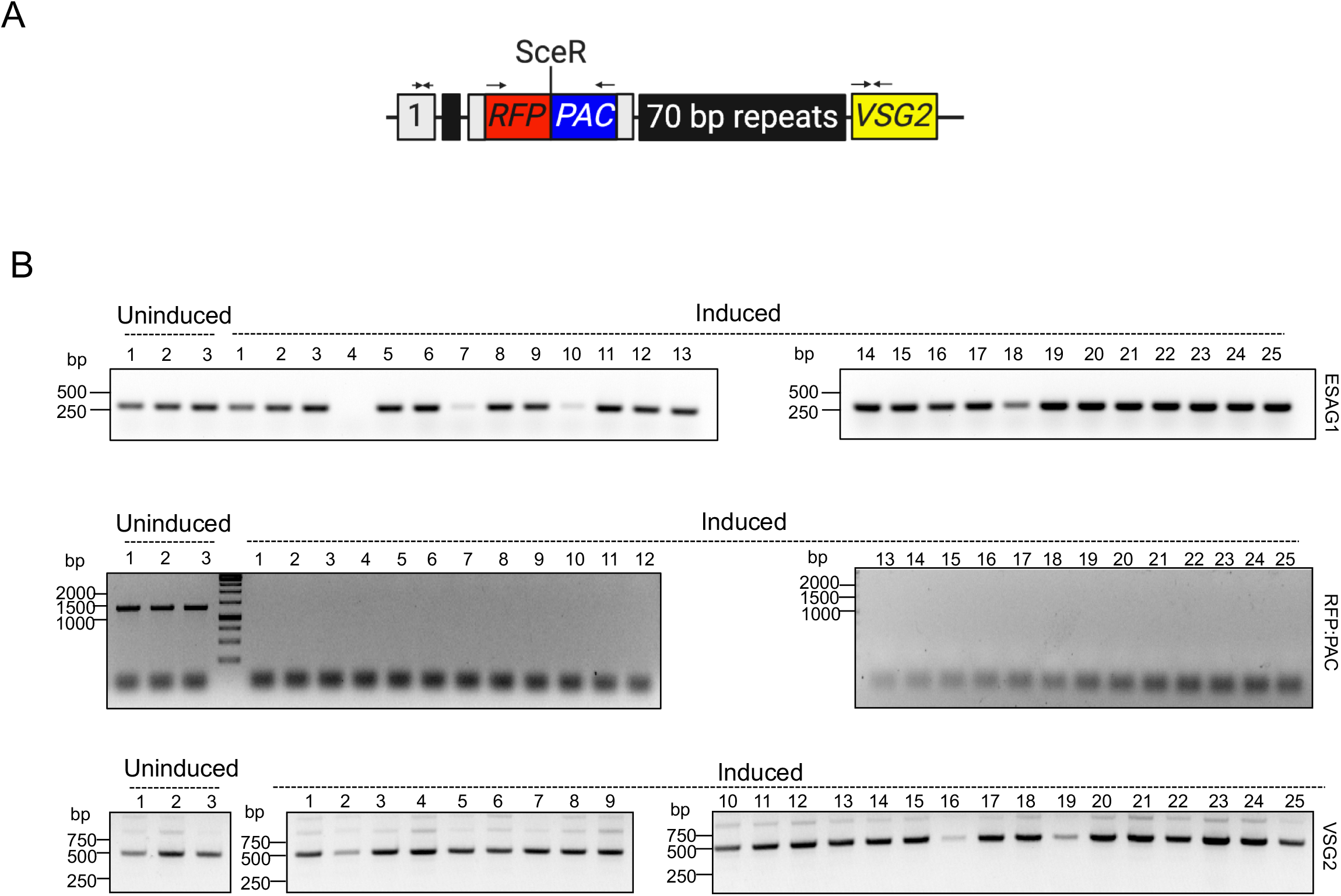
Pseudo^sce^ PCR assays. (A) Schematic indicates the location of the primers used. (B) The PCR assays show the presence or absence of *ESAG1, RFP:PAC* and *VSG2* following an I-SceI induced DSB.

**Supplementary Figure 4:**
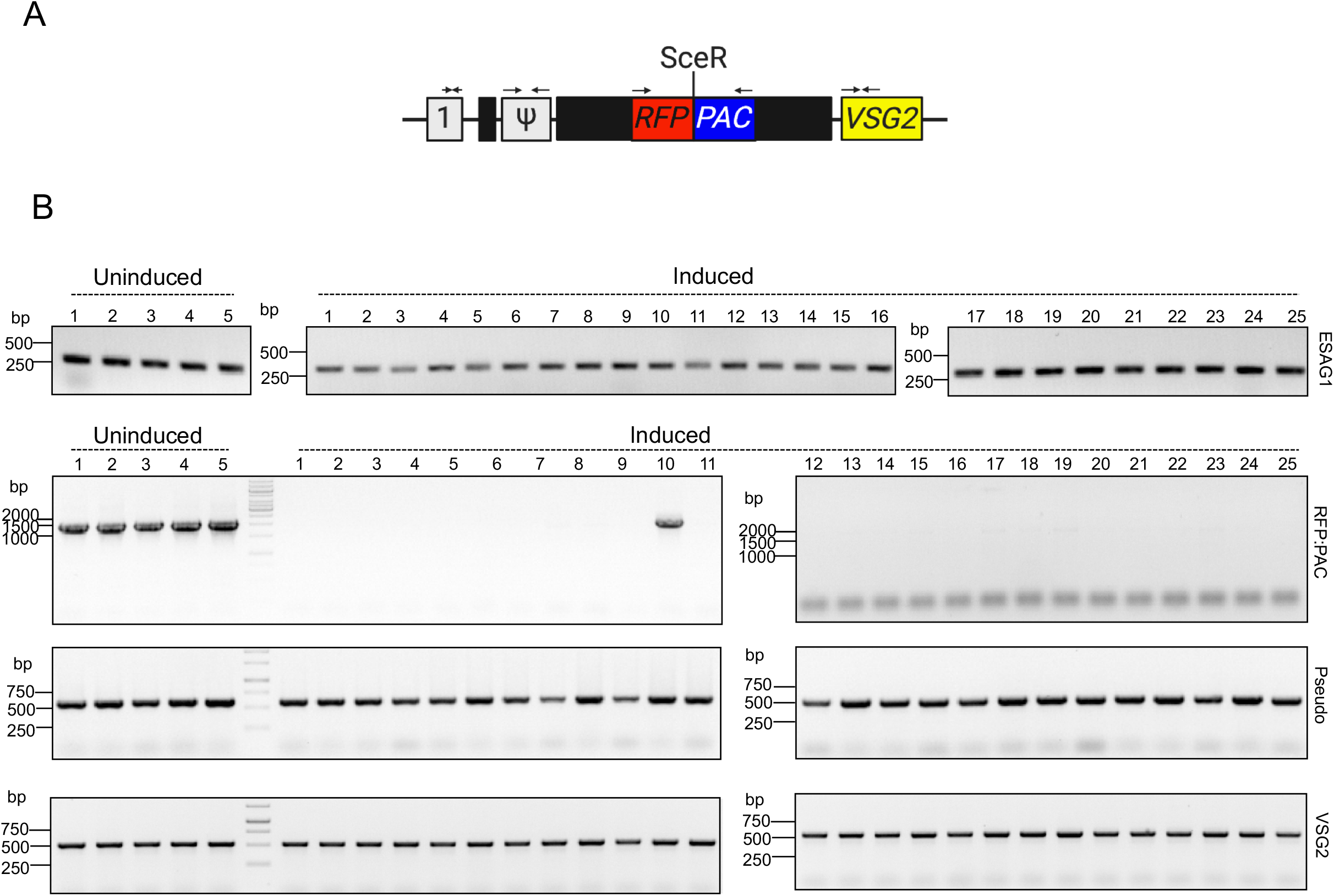
70int^sce^ PCR assays. (A) Schematic indicates the location of the primers used. (B) The PCR assays show the presence or absence of *ESAG1, Pseudo, RFP:PAC* and *VSG2* following an I-SceI induced DSB.

**Supplementary Figure 5:**
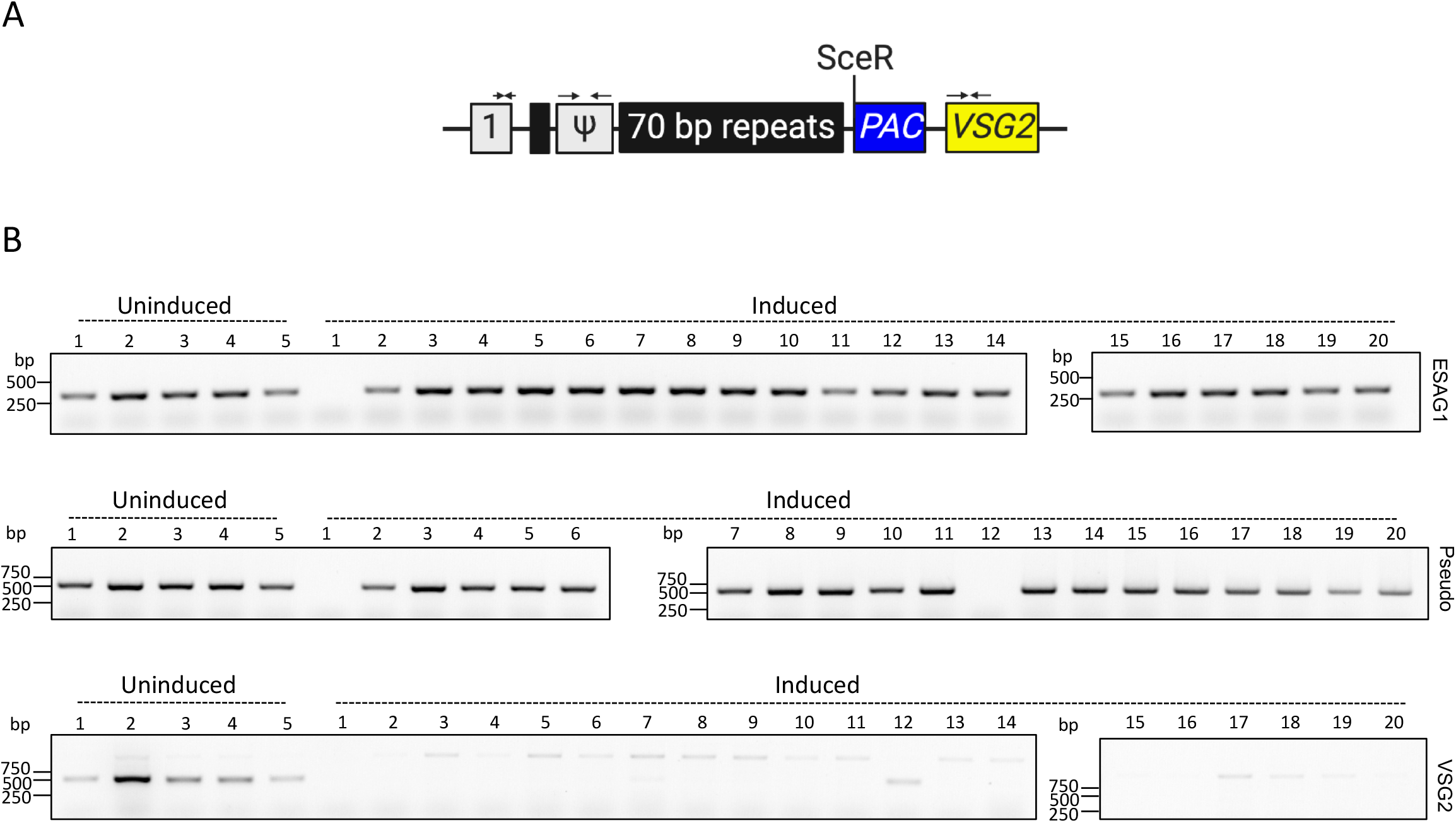
VSG^up^ PCR assays. (A) Schematic indicates the location of the primers used. (B) The PCR assays show the presence or absence of *ESAG1, Pseudo* and *VSG2* following an I-SceI induced DSB.

**Supplementary Figure 6.**
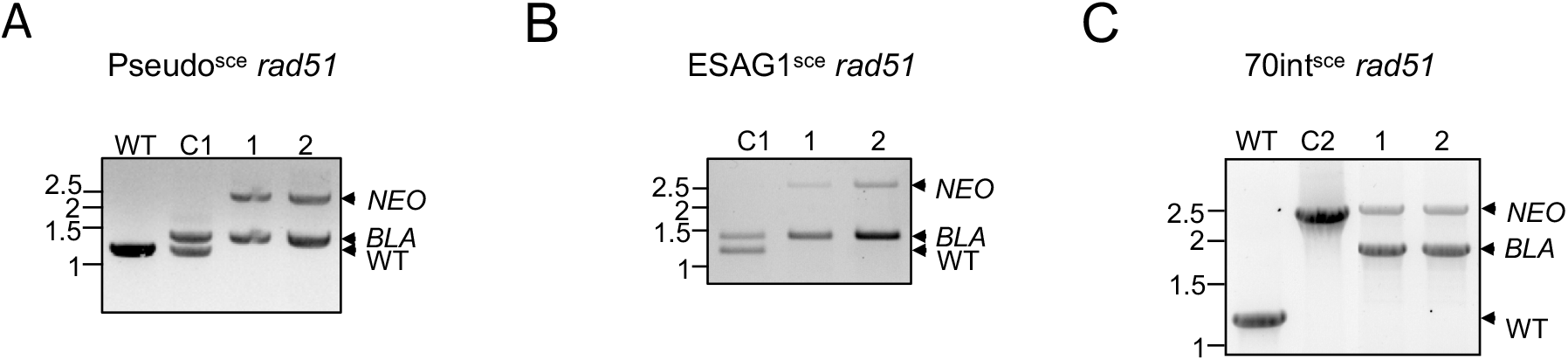
Generation of *rad51* null strains. (A) PCR assay confirming *rad51* double allele replacement Pseudo^sce^. (B) PCR assay confirming *rad51* double allele replacement ESAG1^sce^. (C) PCR assay confirming *rad51* double allele replacement 70int^sce^. *NEO, Neomycin Phosphotransferase; BLA, Blasticidin deaminase*. C1, control plasmid for *BLA;* C2, control plasmid for *NEO*. RAD51 was detected using polyclonal anti-RAD51 antiserum^22^.

**Supplementary Figure 7:**
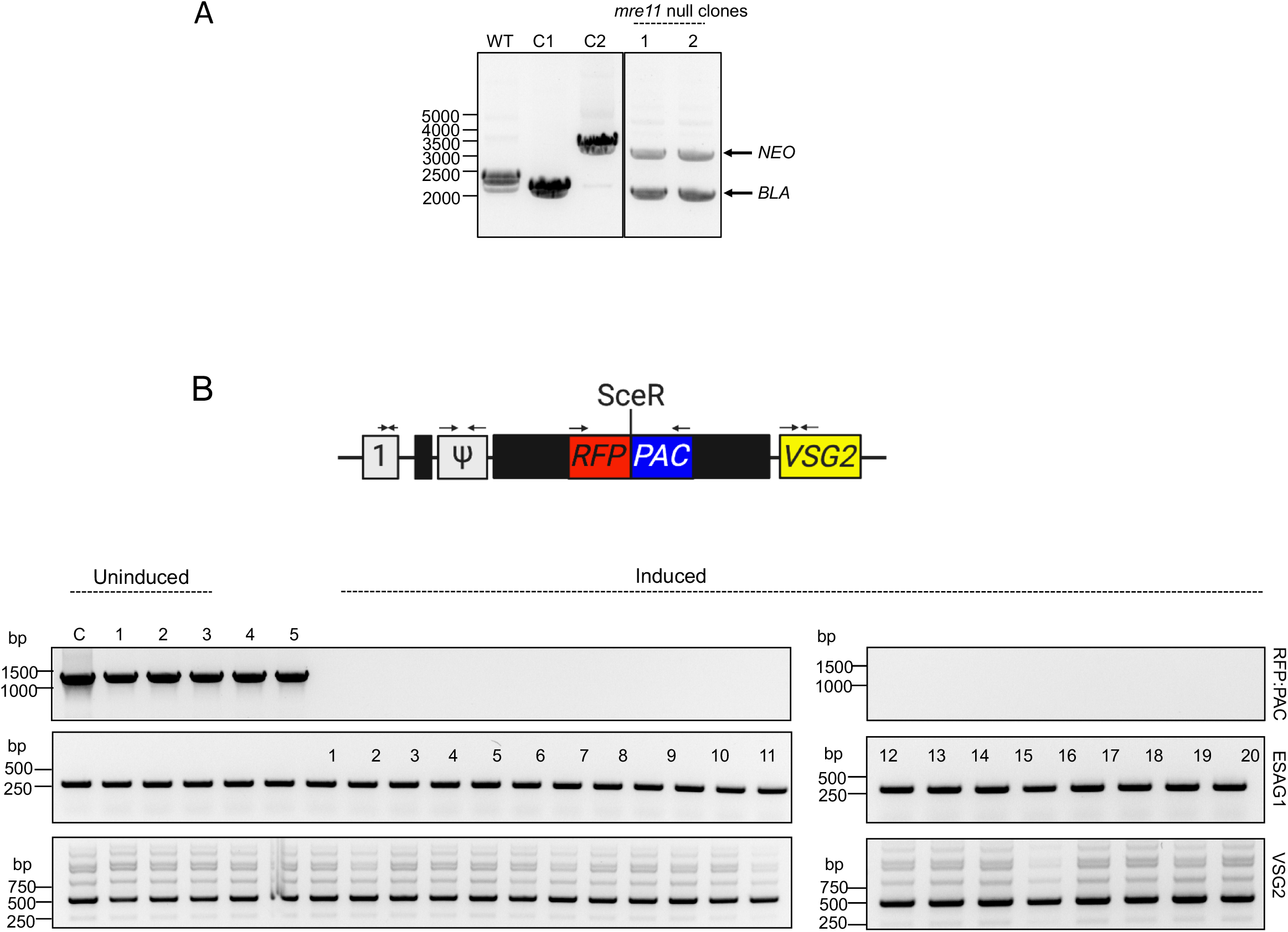
Generation of *mre11* null strains. (A) PCR assay confirming *mre11* double allele replacement. C1, control plasmid for *BLA*; C2, control plasmid for *NEO*. (B) Upper panel: Schematic indicates the location of the primers used. Lower panel: The PCR assays show the presence or absence of *ESAG1, RFP:PAC* and *VSG2* following an I-SceI induced DSB. *NEO, Neomycin Phosphotransferase; BLA, Blasticidin deaminase*.

**Supplementary Figure 8:**
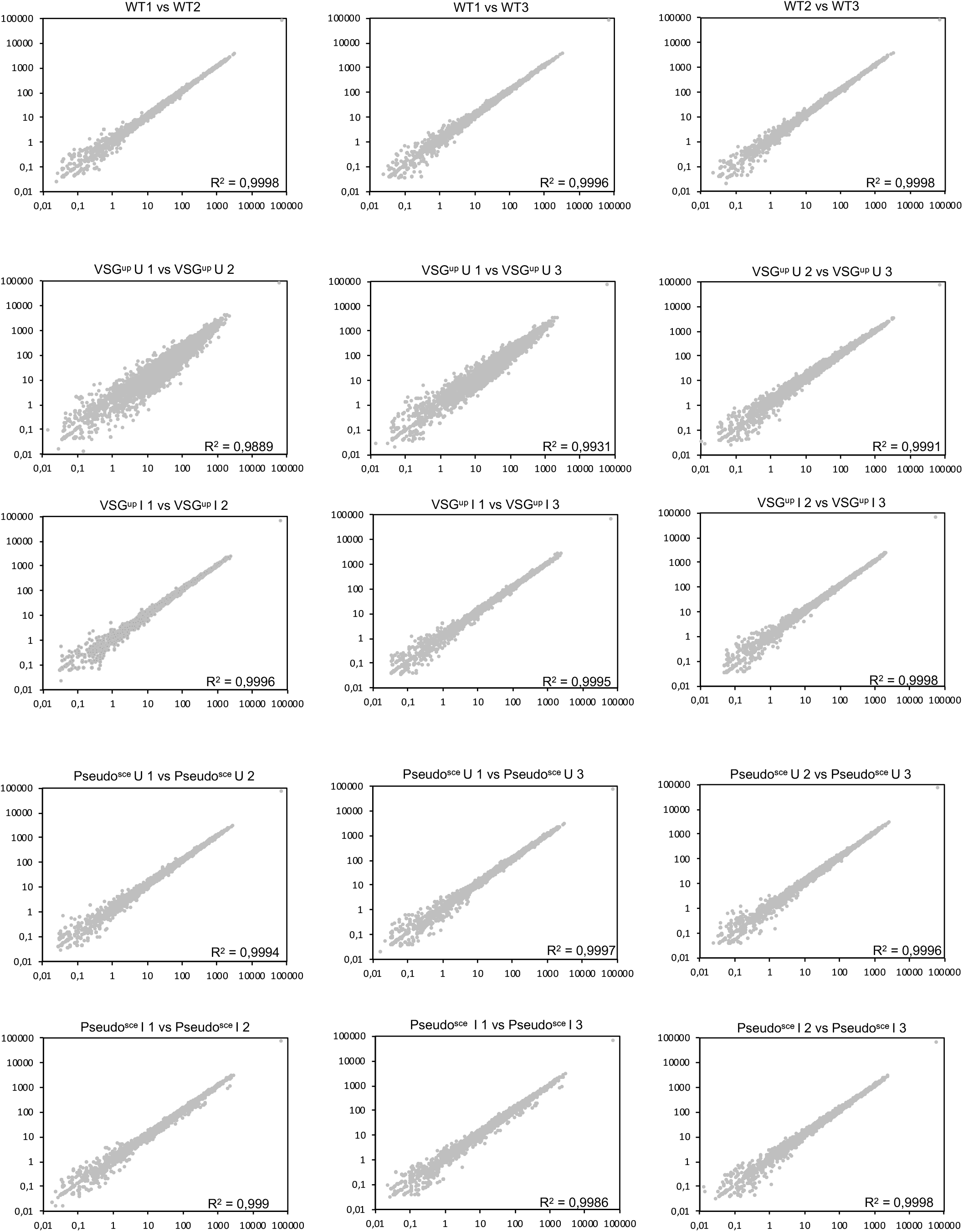
RNA-seq analysis in the VSG^up^ and Pseudo^sce^ cell lines. (A) The scatter plots depict pair-wise comparisons between three biological replicates of wild-type (WT) cells, VSG^up^ uninduced (U) and induced (I) and Pseudo^sce^ uninduced (U) and induced (I).

Table 1: Complete table of all gene expression changes 6 hours following a DSB.

Table 2: Top 100 genes with the greatest fold changes in VSG^up^ and Pseudo^sce^.

## Notes

### Competing Interest Statement

The authors have declared no competing interest.

